# Network architecture producing swing to stance transitions in an insect walking system

**DOI:** 10.1101/2022.01.20.477058

**Authors:** Beck Strohmer, Charalampos Mantziaris, Demos Kynigopoulos, Poramate Manoonpong, Leon Bonde Larsen, Ansgar Büschges

## Abstract

The walking system of the stick insect is one of the most thoroughly described invertebrate systems. We know a lot about the role of sensory input in the control of stepping of a single leg. However, the neuronal organization and connectivity of the central neural networks underlying the rhythmic activation and coordination of leg muscles still remain elusive. It is assumed that these networks can couple in the absence of phasic sensory input due to the observation of spontaneous recurrent patterns (SRPs) of coordinated motor activity equivalent to fictive stepping-phase transitions. Here we sought to quantify the phase of motor activity within SRPs in the isolated and interconnected meso- and metathoracic ganglia. We show that SRPs occur not only in the meso-, but also in the metathoracic ganglia of the stick insect, discovering a qualitative difference between them. We construct a network based on neurophysiological data capable of reproducing the measured SRP phases to investigate this difference. By comparing network output to the biological measurements we confirm the plausibility of the architecture and provide a hypothesis to account for these qualitative differences. The neural architecture we present couples individual central pattern generators to reproduce the fictive stepping-phase transitions observed in deafferented stick insect preparations after pharmacological activation, providing insights into the neural architecture underlying coordinated locomotion.

## 1 Introduction

Walking is based on cyclic patterns of coordinated movement among legs. A step consists of the stance phase, during which the leg has ground contact, supports, and propels the animal, and the swing phase, when the leg is lifted and moves back to its initial position to complete the stepping cycle. We know a lot about the role of sensory organs in initiating or terminating the stepping phases in a single leg (Büschges, 2005). However, the exact organization of the central neural networks underlying stepping-phase transitions in multi-segmented locomotor organs still remains elusive.

So far there have been different approaches considered in modeling studies focusing on either vertebrates or invertebrates. According to one approach, stepping is based on motor neuron synergies controlled by central rhythm and pattern generating networks, which then interact with sensory feedback to generate stepping (McCrea and Rybak, 2008, Markin et al., 2012). However, in other studies sensory feedback is a larger focus, (Ekeberg and Pearson, 2005) reconstruct stepping in a cat’s hindlimb based on sensory signals and the mechanical coupling of the legs. Similarly, in models studying the stick insect walking system, sensory signals play a large role as the position of the leg is the main factor for ending a step phase and initiating the next one (Schilling et al., 2013, Tóth and Daun-Gruhn, 2016). Lastly, there are reduced models which define a single network for each leg and do not consider the individual networks controlling each leg segment (Holmes et al., 2006, Schilling et al., 2013).

For our modeling approach we have decided to use biological data from the stick insect because it is known that there are dedicated central pattern generators (CPGs) to move each of the main leg joints (Büschges et al., 1995). Walking in the stick insect is based on intra- and intersegmental interaction among CPG networks and peripheral sensory input (Bidaye et al., 2018). The relative contribution of the central versus peripheral neuronal mechanisms for walking has been often debated. Pharmacological activation of the central networks in deafferented insect preparations has contributed significantly to our understanding regarding the possible role of sensory input in adapting the default centrally-generated patterns, potentially giving rise to behaviorally-relevant coordination. Such experiments have not resulted in generation of motor patterns similar to the coordination patterns observed *in vivo*. Thus, there has been no indication of fictive walking in adult insect preparations (Mantziaris et al., 2017, Knebel et al., 2017, Mantziaris et al., 2020). However, previous experiments in the deafferented stick insect mesothoracic ganglion, after application of the muscarinic agonist pilocarpine, have indeed revealed coordinated activity of the motor neuron pools within a hemisegment in the absence of phasic sensory input. This activity resembles stepping-phase transitions *in vivo* (Büschges et al., 1995) and are therefore referred to as fictive stepping-phase transitions. There have been distinct patterns of coordinated motor activity described which spontaneously emerge in a non-cycle-to-cycle fashion throughout the recording, and are therefore called spontaneous recurrent patterns (SRPs). The SRP1 is characterized by a switch from protractor to retractor motor neuron activity during a depressor burst consisting of both the slow and fast depressor motor neurons. This switch resembles a transition from swing to stance stepping phase during forward walking. The SRP2 is characterized by the opposite switch from retractor to protractor activity during a combined slow and fast depressor burst, resembling a transition from swing to stance stepping phase during backward walking (Büschges et al., 1995, Büschges, 1995). These patterns have only been described for the mesothoracic ganglion of the stick insect so far, with the SRP1 showing a higher occurrence throughout the recording compared to SRP2. Although there have been neurons identified in the ventral nerve cord which influence the occurrence of SRPs, the underlying mechanisms resulting in SRP generation are still unknown (Mantziaris et al., 2020, Büschges, 1995).

SRP-like activity patterns have also been reported in neurophysiological recordings of motor activity in the fourth thoracic ganglion of the crayfish after bath application of cholinergic agonists (Chrachri and Clarac, 1990). In their study, 90% of the recorded intervals showed coordinated activity consisting of the levator motor neurons bursting in phase with the retractor motor neurons. This pattern is indicative of fictive backward walking. In the stick insect mesothoracic ganglion, according to the only study focusing on intrasegmental coordination in insects, 78% of the patterns of coordinated activity belonged to the SRP1 type, representing a fictive transition from swing to stance phase during forward stepping (Büschges et al., 1995). Here, we hypothesize that SRPs can also be found in other ganglia, such as the metathoracic ganglion, and that an underlying network architecture exists to generate these SRPs in the absence of sensory feedback. We have chosen to investigate potential architectures through simulation, following the sentiment by (Grillner, 2003) that “… many interactive processes on the subcellular, cellular and network levels are dynamic and complex. Computational methods are therefore required to test whether tentative explanations derived by intuition can account for experimental findings.”

The majority of current engineering research develops coordination through feedback and subsequent rules based on the returned values. Popular controllers such as WalkNet (Schilling et al., 2013) and neuroWalknet (Schilling and Cruse, 2020) use behavioral data from biology to create an artificial neural network capable of controlling 18 degrees of freedom. The individual leg controllers use coordination rules, network architecture, and sensory feedback to produce walking behaviors on a six-legged robot. WalkNet can perform forwards and backwards walking on smooth and uneven terrain as well as curved walking. neuroWalknet extends the walking ability to produce different footfall patterns including tripod, tetrapod, and pentapod patterns as well as other stable intermediate patterns observed in stick insects (Graham, 1972) and Drosophila (Wosnitza et al., 2013). Ekeberg et al. (2004) realize a neuronal control network sufficient for controlling stick insect legs in each segment - front, middle, and hind. They use a simulation to send feedback to the control network as the primary driver for joint coordination. von Twickel et al. (2011) use a similar strategy by decentralizing the joint controllers and relying on feedback from a simulated leg to develop a single-leg controller. All of the mentioned studies have proposed different control models for joint (intralimb) and leg (interlimb) coordination to reproduce biological behaviors and confirm the importance of sensory feedback in coordinated walking. However, there have not been any studies investigating the underlying neural network architecture before sensory feedback is applied in order to reproduce the fictive stepping-phase transitions observed in deafferented stick insect preparations. We address this by using biologically-plausible spiking and non-spiking neuron models to create a single joint architecture similar to the one suggested by Yeldesbay and Daun (2020). The setup connects spiking rhythm-generating popluations (RGPs) to non-spiking interneurons (NSIs) to inhibit the spiking motor neuron populations (MNPs). Coordination NSIs (cNSIs) are also used to connect the individual joints to create inter-joint coordination similar to the biological interneurons found by Büschges (1995). The addition of NSIs is able to decouple the neuronal dynamics of the spiking populations, removing the issue of competing dynamics when communicating between spiking neurons. In this way, we find that the integration of NSI’s into spiking neural networks reduces complexity when building the network while maintaining biological plausibility.

Our study shows that SRPs occur in both the meso- and metathoracic ganglia of the stick insect and highlight an interesting difference between the two thoracic ganglia. We propose a neural architecture that is able to reproduce the prominent SRP in each ganglia confirming that network architecture can account for the difference in SRP type.

## 2 Methods

### 2.1 Neurophysiology

#### 2.1.1 Experimental Animals

The animals used in this study are adult female Indian stick insects of the species *Carausius morosus* bred in our colony at the Biocenter, University of Cologne. Animals are kept at 20 to 26°C with 45 to 60% humidity, under a 12 hour light/12 hour dark cycle. The experimental procedures described below comply with the German National and State Regulations for Animal Welfare and Animal Experiments.

#### 2.1.2 Preparation and experimental setup

The experimental procedure followed in this study has been previously established by Büschges et al. (1995). Some experiments are performed on the ipsilateral hemisegment of the isolated (disconnected from neighboring ganglia) meso- and metathoracic ganglia. Other experiments are performed on the interconnected meso- and metathoracic ganglia. In all studies, the ganglia are deafferented and isolated from the rest of the nerve cord. Additionally, the first abdominal ganglion is always left interconnected to the metathoracic ganglion. Rhythmic activity in motor neuron pools is induced by bath application of the muscarinic acetylcholine receptor agonist pilocarpine, and is assessed by recording extracellular activity by placing extracellular electrodes on the lateral nerves nl2, nl3, nl5, and C2. This captures the motor activity innervating the leg muscles. All lateral nerves in the ganglia of interest are either crushed or cut to block afferent input. The nerves nl2 and nl5 carry the axons that innervate the antagonistic protractor and retractor coxae muscles, while nl3 and C2 innervate the extensor tibiae and depressor trocanteris respectively (Goldammer and Schmidt, 2012). The protractor and retractor move the leg forwards and backwards along the horizontal plane, the extensor muscle extends the tibia and the depressor muscle allows for downward movement of the leg along the vertical plane. Collectively, these muscles allow the movement about the three main leg joints: the Thorax-Coxa (ThC), the Coxa-Trochanter (CTr) and the Femur-Tibia (FTi) joint.

The signal recorded with the extracellular electrodes is pre-amplified by 100-fold using isolated low-noise preamplifiers (model PA101, Electronics workshop, Zoological Institute, Cologne). It is further amplified by ten-fold to reach an overall gain of 1000 and filtered (low-cut: 200Hz, high-cut: 3kHz) using a standard 4-channel amplifier/signal conditioner (model MA102, Electronics workshop, Zoological Institute, Cologne). Finally, the signal is digitized at a sampling rate of 12kHz, using the Micro 1401-3 acquisition unit (CED, Cambridge, UK) and it is monitored using the Spike2 software (CED, Cambridge, UK).

#### 2.1.3 Analysis of the neurophysiological data

The electrophysiological data are initially assessed by using tools provided by Spike2. Time series of the action potentials (spikes) are marked by manually setting a threshold. In cases where spikes of non-interesting neurons also cross the threshold, such as those of the common inhibitory motor neuron, the respective time-series are subtracted from the data. The depressor bursts are marked, noting the timing of the burst onset because the depressor cycles are used as a reference for the analysis.

Fig 1 depicts how the phase difference between nerves is calculated. In this study, “Nerve 2” in Fig 1 represents bursts of motor activity in the depressor because it is the reference nerve. Spontaneous recurrent patterns of coordinated motor activity are manually marked throughout the recording to use for the phase calculation. The spike time-series are exported using a sampling rate of 1000Hz and the phase of each spike within the ongoing depressor cycle is calculated throughout the recording using MATLAB. Phase values are binned (bin size = 10°) and the mean bin value (± standard deviation (STD)) among animal preparations is plotted after normalizing data of each animal to the maximum number of spikes. Histograms are made by either considering all depressor cycles throughout the recording or selecting only those cycles during which an SRP1 or SRP2 occurred. Circular means and angular deviations, as well as the 90% confidence intervals (CI) are calculated using the Circular Statistics Toolbox in MATLAB (Berens, 2009).

**Figure 1:**
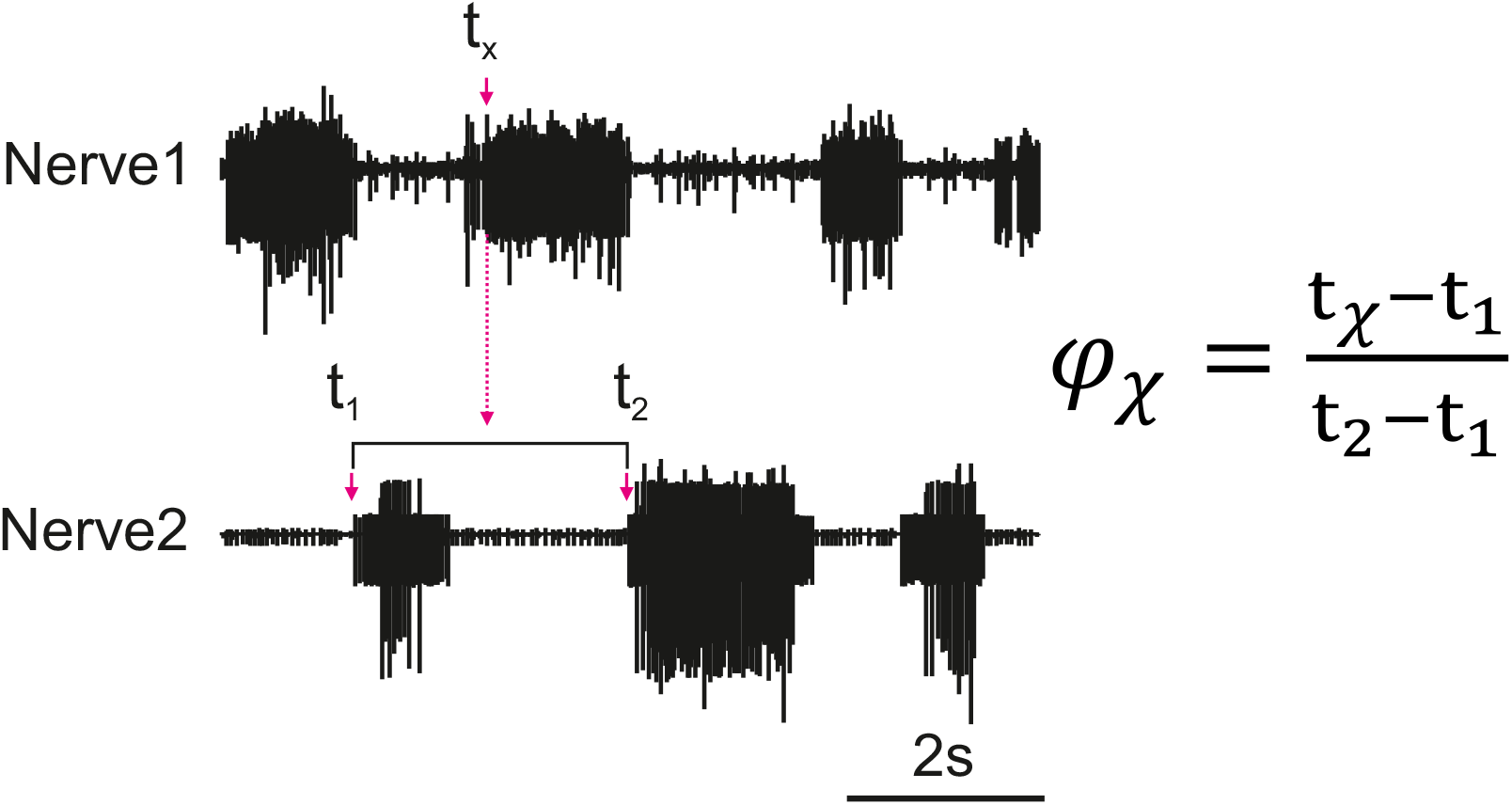
Illustration of how the phase (*ϕ*_*x*_) of each spike of a nerve (Nerve 1) is calculated in relation to the cycle of a reference nerve (Nerve 2). The time is recorded for all spikes reaching the set threshold within Nerve 1 (*t*_*x*_). The phase is calculated for each spike and then averaged to find the average activity of a nerve as compared to the reference nerve. The cycle period is dynamic so the phase is calculated based on the corresponding cycle, the cycle period is represented by *t*_2_ − *t*_1_ where *t*_1_ is the cycle onset and *t*_2_ is the end of the cycle.

### 2.2 Network Simulation

The neural network designed in this study is based on observations from biological research. Individual CPGs are known to control a single antagonistic muscle pair producing alternating rhythmic activity per joint (Büschges et al., 1995, Bidaye et al., 2018). This means that the individual networks must be coordinated to produce the observed swing and stance phases of a single leg step cycle. Membrane potential oscillations of NSIs have been shown to correlate with SRPs (Büschges, 1995) indicating that NSIs are most likely involved in inter-joint coordination. Our network uses this knowledge to couple three CPG networks with NSIs to produce coordinated firing.

#### 2.2.1 Single Joint Architecture

Before coordination can be achieved, a reliable anti-phasic output must be generated corresponding to the antagonistic muscle pairs in the stick insect leg controlling each joint. Each individual joint is controlled by one CPG consisting of two RGPs mutually inhibiting each other in a half-center oscillator architecture (Bidaye et al., 2018). Fig 2a shows a detailed diagram of the neural network controlling a single joint.

**Figure 2:**
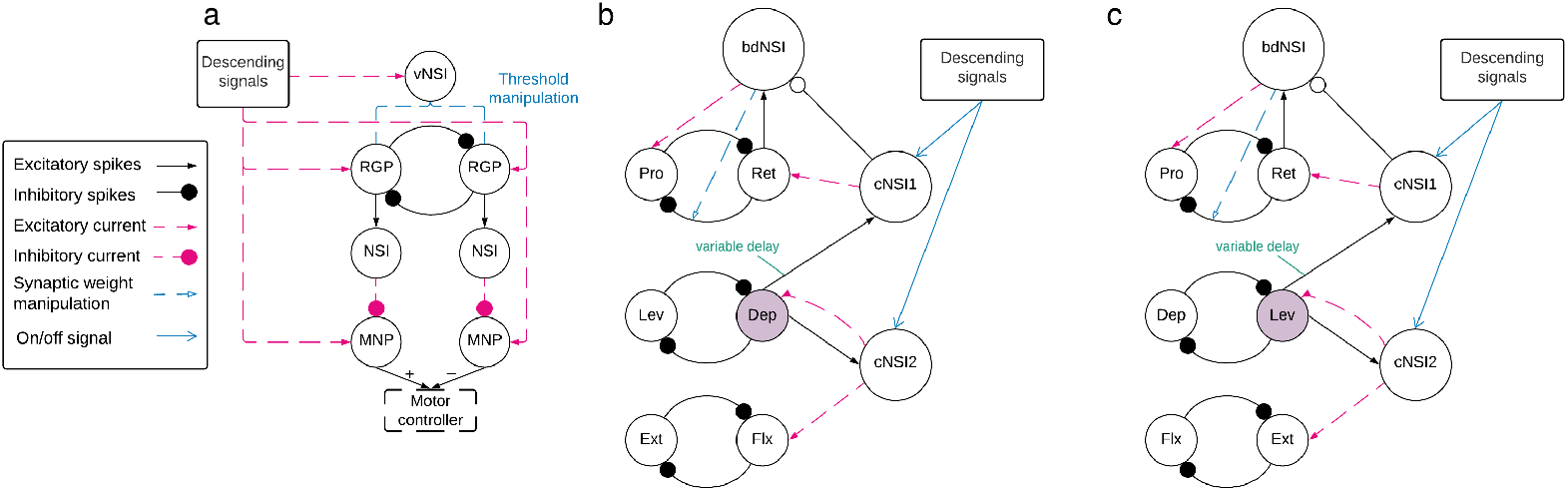
**a**. Network architecture of an individual joint showing the half-center oscillator comprised of two RGPs mutually inhibiting each other with a synaptic weight of − 5000*nS*. The RGPs excite buffer NSIs which then inhibit the MNPs periodically, creating output oscillations. The voltage threshold potential *V*_*th*_ of neurons in the RGPs is updated based on the input current to the velocity NSI (vNSI) in order to manipulate output frequency. Excitatory current is sent to the RGPs to initiate bursting and to the MNPs to promote tonic spiking. The output of the MNPs is sent to a motor controller. **b and c**. The network architecture for coupling individual joints through coordination NSIs (cNSIs). The rhythm-generating populations are labeled based on the MNPs they control - Protractor (Pro), Retractor (Ret), Depressor (Dep), Levator (Lev), Extensor (Ext), and Flexor (Flx). The highlighted populations are the RGPs driving coordination. The burst duration NSI (bdNSI) is used during uncoordinated firing and silenced when the cNSIs are active. The retractor RGP sends spikes to the bdNSI. When the bdNSI is active, it waits a calculated amount of time before sending excitatory current to the protractor RGP and reducing the inhibition from the retractor to protractor RGP in order to stop the Pro MNP from bursting. **b**. The architecture to produce SRP1 transitions. The Dep RGP drives coordination, sending excitatory spikes to the cNSIs to coordinate bursting through excitatory current to the Flx and Ret RGPs. **c**. The architecture to produce SRP2 transitions. The Lev RGP drives coordination, sending excitatory spikes to the cNSIs to coordinate bursting through excitatory current to the Ext and Ret RGPs.

The RGPs and MNPs consist of adaptive exponential integrate-and-fire (AdEx) neurons. The parameters for the RGP neurons are set to bursting whereas the parameters used for the MNP neurons produce tonic spiking. These parameters are set according to (Naud et al., 2008). The AdEx neuron dynamics are described by Eqs (1) and (2).

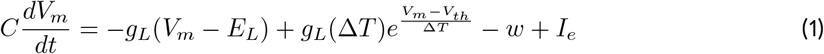

when *V*_*m*_ > 0*mV* then *V*_*m*_ → *V*_*reset*_

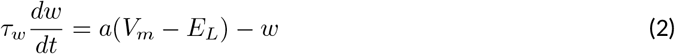

when *V*_*m*_ > 0*mV* then *w* → *w* + *b*

“where C is the membrane capacitance, *V*_*m*_ is the membrane potential, *E*_*L*_ is the resting potential, *g*_*L*_ is the leakage conductance, *I*_*e*_ is the bias current plus Gaussian white noise, *a* is the sub-threshold adaptation conductance, *b* is the spike-triggered adaptation, Δ_*T*_ is the sharpness factor, *τ*_*w*_ is the adaptation time constant, *V*_*th*_ is the voltage threshold potential, *V*_*reset*_ is the reset potential, and *w* is the spike adaptation current (Naud et al., 2008). Eq (1) defines the change in membrane potential per time step whereas Eq (2) outlines the current adaptation.” (Strohmer et al., 2021)

All NSIs used within the network are modeled as leaky integrate-and-fire neurons with a high enough voltage threshold to avoid spiking. The neuronal dynamics are described by Eq (3).

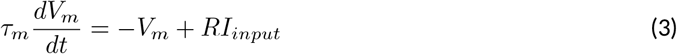

“where *τ*_*m*_ = *RC* is the membrane time constant, *V*_*m*_ is membrane potential, *I*_*input*_ is input bias current plus Gaussian white noise, and *R* is membrane resistance.” (Strohmer et al., 2021)

The RGPs are initialized with an excitatory current of 500*pA* but this switches to a frequency regulating excitatory current. The current is designed to be low enough to induce bursting while producing the desired frequency. This current increases with frequency according to Eq (4).

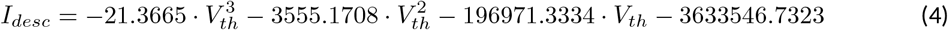

Where *I*_*desc*_ is the descending input current to the RGPs and *V*_*th*_ is the voltage threshold potential. The parameter calculations defined within the network architecture depend on frequency. In previous work, Strohmer et al. (2020) found a linear relationship between *V*_*th*_ and frequency. Therefore, all linear relationship calculations use *V*_*th*_ as a substitute to approximate frequency instead of calculating frequency at each time step. This substitution reduces computational load.

The descending signals act as an “on/off” switch for the RGPs. Descending signals are also able to control the frequency of output oscillations from the RGPs by sending excitatory current to the velocity NSI (vNSI). This setup is based on biology, Berendes et al. (2016) suggest that there could be multiple pathways for descending signals and that the signals controlling output frequency are connected to CPGs through sensory neurons. Validation that the vNSI is able to control output frequency by manipulating the *V*_*th*_ of the RGP neurons was previously shown by Strohmer et al. (2021). The stick insect steps at a frequency of approximately 1 − 4*Hz* on a slippery surface (Graham and Cruse, 1981). These observed speeds are used to constrain the output frequency per joint during testing. Our work investigates the phase relationships observed in biological recordings of low-frequency motor activity in deafferented preparations to ultimately reproduce the phase relationships observed during live stepping. In this way, our aim is to match the phase relationship measured from deafferented samples and then visually confirm if this produces stepping-phase transitions *in vivo*. We can achieve this confirmation using a robot leg in simulation. We are able to directly compare network simulation and biological measurements by normalizing over degrees of a step cycle instead of timing, thereby, removing the problem of the difference in step cycle period.

In our network, the descending signals also send an excitatory current of 500*pA* to the MNPs to allow the motor neurons to begin tonically spiking. The rhythm of the network’s output is finally determined by phasic inhibition. From the RGPs, excitatory spikes are sent to the buffer NSIs. These NSIs act to inhibit the MNPs, mimicking the network of the stick insect (Yeldesbay and Daun, 2020, Mantziaris et al., 2020). The synaptic weight from the RGP determines the extent of membrane potential fluctuation of the NSI. We limit the fluctuation to approximately 15*mV* to keep within a biologically-plausible range (Burrows and Siegler, 1978). The dependence of synaptic weight on frequency (*V*_*th*_) is outlined in Eq (5).

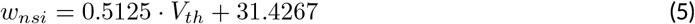

Where *w*_*nsi*_ is the synaptic weight from the RGPs to the NSIs and *V*_*th*_ is the voltage threshold potential of the AdEx neurons in the RGP population.

The inhibitory current from the NSIs to the MNPs is only dependent upon the momentary membrane potential of the NSI. This is described in Eq (6).

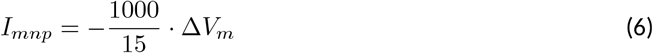

Where *I*_*mnp*_ is the current from the NSI to the MNP and *V*_*m*_ is the membrane potential of the NSI. The selection of 15 as the denominator ensures that the maximum inhibition is −1000*pA*. The maximum is determined through trial and error.

#### 2.2.2 Interjoint Coordination

Fig 2b and c show a high level overview of the neural network for coordinating firing between joints. Each of these sub-figures represents the same neuronal populations, it is only split so that the relevant synaptic connections can be highlighted based on the desired SRP output. Each joint is represented by a single half center oscillator to increase the clarity of the figure even though the controller per joint is as shown in Fig 2a. Network coordination is achieved through the use of cNSIs which receive excitatory spikes from the depressor or levator RGP depending on walking direction. The synaptic weight from the RGP to the cNSI is determined by Eq (7).

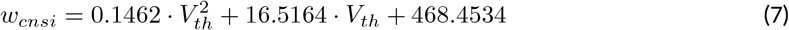

Where *w*_*cnsi*_ is the synaptic weight from the pre-synaptic RGPs to the cNSI and *V*_*th*_ is the voltage threshold potential of the AdEx neurons in the RGPs. The weight is determined through trials and ensures the cNSI’s membrane potential fluctuates by a maximum of 11*mV* as is observed in biological NSIs found within networks generating SRPs (Büschges, 1995). It should be noted that this change in membrane potential is smaller than the generic buffer NSIs used in the single joint architecture.

The cNSI connects to the post-synaptic RGPs with a weight determined by Eq (8). During coordinated firing, the cNSIs are selected as active by the descending signals, allowing current to pass through them and coupling the joints.

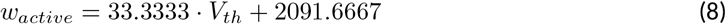

Where *w*_*active*_ is the synaptic weight from the cNSIs to the post-synaptic RGPs and *V*_*th*_ is the voltage threshold potential of the AdEx neurons in the RGPs. In order to send current from a cNSI to a post-synaptic RGP, the weight is multiplied by the change in membrane potential of the cNSI at each time step as shown in Eq (9).

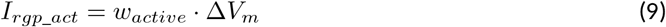

Where *I*_*rgp*_*act*_ is the current from the cNSI to the post-synaptic RGP and *V*_*m*_ is the membrane potential of the cNSI.

During coordinated firing for the SRP1 swing to stance transition, the cNSIs are selected as active and Eq (9) determines the excitatory current sent to the Flx and Ret RGPs. During SRP2 coordination, the cNSIs send excitatory current to the Ext and Ret RGPs. The current injections from the cNSIs drive coordination between each of the joints but do not necessarily produce the same phase differences as recorded in deafferented samples. In order to tune the phase difference between joints, the synaptic delay from the CTr joint RGP to cNSI1 is modified (see synapse marked with “variable delay” in Fig 2 b, c). The other synaptic delays are held constant and can be seen in Table 1.

**Table 1:** Synaptic delays between spiking populations and NSIs.

| Synapse | Synaptic Delay (ms) |
| --- | --- |
| CTr RGP to cNSI1 | variable |
| CTr RGP to cNSI2 | 2 |
| ThC RGPs to buffer NSIs | 5 |
| CTr RGPs to buffer NSIs | 5 |
| FTi RGPs to buffer NSIs | 2 |

Finally, Fig 2 b, c show feedback current from cNSI2 to the driving CTr RGP. This feedback between the FTi and CTr joint has been observed in biological networks (Daun-Gruhn and Büschges, 2011) and is added to our network to increase biological fidelity. The strength of the feedback current is set as 0.1% of the current from the cNSI (Eq (9)). The strength of this current is determined through manual testing.

#### 2.2.3 Protractor Burst Duration Modulation

Deafferented preparations of the stick insect indicate a decoupling of protraction burst duration from walking frequency during uncoordinated firing. Büschges et al. (1995)’s deafferented sample recordings show that retractor burst duration is correlated to cycle period but protractor burst duration is not. Therefore, this phenomenon must be accounted for when simulating uncoordinated firing to maintain biological plausiblity. We were able to reproduce the decoupling of protractor burst duration from cycle period using a burst duration NSI (bdNSI). The architecture is shown in Fig 2b and c. The presented solution is inspired by biological studies that find that blocking of calcium-dependent ion channels is able to lengthen burst duration of a neuron (Grillner, 2003). We replicate this by increasing current to the protractor RGP and reducing inhibition from the retractor RGP. The prolonged excitation of the protractor RGP inhibits the protractor MNP, reducing the burst duration to the muscle. Implemented in practice, this means the retractor RGP sends excitatory spikes to a bdNSI. When the retractor RGP is spiking, the protractor MNP is also spiking so these excitatory spikes serve to inform the system that the protractor MNP has started spiking. Once the bdNSI’s membrane potential is depolarized by 2*mV*, the system waits for a period of time based on the frequency of stepping before sending an excitatory current of 3000*pA* to the protractor RGP and reducing inhibition from − 5000*nS* to 0*nS* from the retractor to protractor RGP. The waiting time is defined by Eq (10).

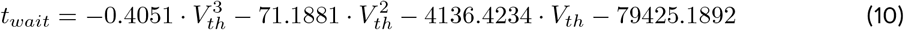

Where *t*_*wait*_ is the amount of time in milliseconds that the protractor RGP is inhibited by the retractor RGP thus allowing the protractor MNP to spike. *V*_*th*_ is the voltage threshold potential of the AdEx neurons in the RGPs. As the cycle period increases due to the frequency slowing down, the amount of time that the protractor MNP is allowed to spike is reduced. The length of inhibition time to the protractor RGP is found through testing.

This mechanism is added to the network architecture and allowed to modulate the burst duration during uncoordinated firing. When the cNSIs are active to generate coordinated firing between joints, the bdNSI is inhibited so that burst duration is not modulated. The plot showing protractor burst duration remaining constant over increasing cycle period during uncoordinated firing is shown in Fig S1 in the Supplementary Material.

Equations (4), (5), (7), (8), and (10) are found through manual testing, plotting the values, and using the polynomial fit function in Python.

#### 2.2.4 Testing

The network architecture is confirmed by comparing simulation output to measurements from the deafferented stick insect. The phase data is extracted from each and equated by normalizing the timing into a 360 degree step cycle.

Fig 3 is a visual illustration of how the phase difference and burst duration are calculated from the simulation results. The calculation of biological phase difference is outlined in the Neurophysiology section (Fig 1).

**Figure 3:**
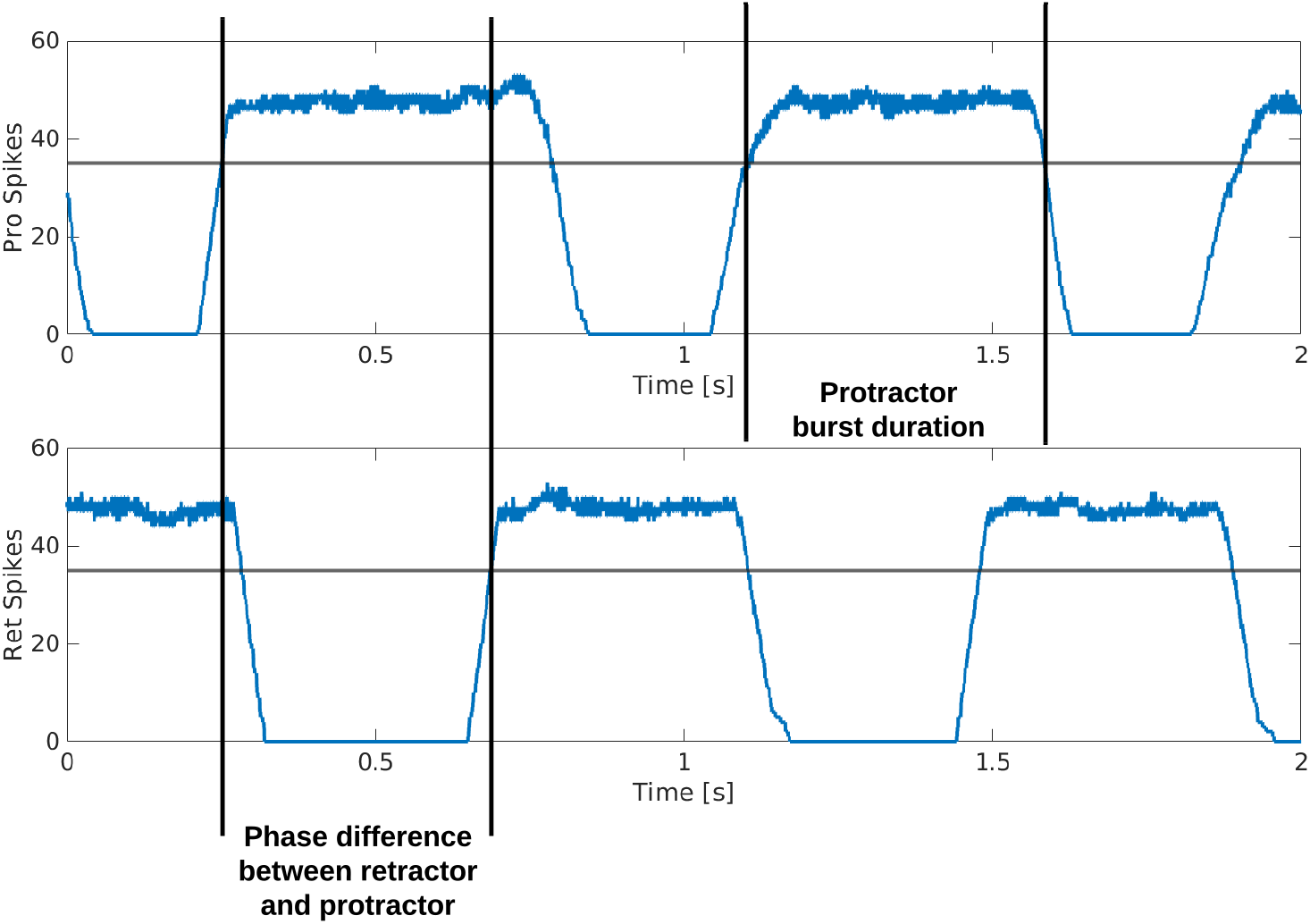
Example output from the protractor and retractor MNPs over two seconds of simulation. A sliding time window of 5 ms is used to count the number of spikes per step. The phase difference is calculated by subtracting the time one MNP crosses a threshold from when the compared MNP crosses the same threshold. This time is divided by the cycle period and multiplied by 360 to produce the phase difference in degrees. The burst duration is calculated by subtracting when an MNP crosses the threshold with a positive slope from when the same MNP crosses the threshold with a negative slope.

The output from each MNP is rate-coded using a sliding time window of 5ms. At each step, the number of spikes are counted within the time window and plotted as the y-value, the time window then moves by the time resolution of the simulation, 0.1ms, and the spikes are counted again to plot the next point.

Each simulation is run for an equivalent of 8 seconds using the Neural Simulation Tool (NEST) (Jordan et al., 2019) (source code available on GitLab (Strohmer, 2021)). The rate-coded spike data is saved and analyzed in MATLAB. Fig 3 shows how the calculations are made based on the simulation output. A threshold of 35 spikes is used because it removes noise from the data. When comparing phase difference of different MNPs, the time step for each MNP as it crosses the 35 spike threshold with a positive slope is used. The time step is converted to actual time 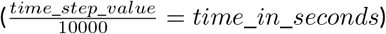 and the values are subtracted and normalized to find phase in degrees, as shown in Eq (11).

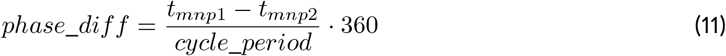

Where *phase_diff* is the phase difference in degrees between the MNPs bursting and *t*_*mnp*1_, *t*_*mnp*1_ are the times in seconds when each MNP crosses the threshold. The cycle period is determined by averaging the period over a single simulation during coordinated firing. This is used to calculate the individual phase differences between MNP bursts. The single phase difference value presented in the results is the average of all these calculations (Eq (11)) during coordinated firing of a single simulation. There exists biological variability between animals as high as 32% in the most extreme case when comparing the phase difference of the Ret-Dep motor activity between observed Animals 1 and 2 (Table S1 in the Supplementary Material). Therefore, we have allowed a buffer of 20°to either side of the circular mean ± angular deviation of a single animal (equivalent to 11%). This is increased to 40°(equivalent to 22%) if the angular deviation was less than 10°. This buffer allows a variability of 11-22%, keeping our simulation within the variation observed between animals.

The burst duration is calculated by looking at a single MNP’s rate-coded output and noting when it crosses the threshold as the slope is increasing and decreasing. This calculation is not normalized, it subtracts the time steps and divides by 10, 000 to formulate the burst duration time in seconds. When testing burst duration, uncoordinated firing is allowed for the complete simulation time of 8 seconds and the presented burst duration time is an average of all phase difference calculations from a single simulation.

Noise is added to the network through current noise to the RGP and MNP neurons. The noise is centered around 0mV with increasing amounts of standard deviation. All tests are run at 5 noise levels, starting at 100pA of current noise and increasing to 500pA by increments of 100pA. The standard deviation of noise to all NSIs in the architecture is held constant at 25pA.

## 3 Results

### 3.1 Neurophysiology

#### 3.1.1 Intrasegmental CPGs can be coordinated in the deafferented preparation

Extracellular recordings of the ipsilateral motor activity in the isolated and deafferented mesothoracic ganglion after pharmacological activation with pilocarpine reveal patterns of coordinated activity similar to the SRPs previously reported by Büschges et al. (1995) (Fig 4a). SRP1 and SRP2 are denoted in Fig 4a1 and Fig 4a2 with a dashed magenta or orange line respectively. The lines demarcate the onset of the respective depressor burst. All highlighted patterns on Fig 4a1 belong to the SRP1 type. Each time there is combined activity of both the fast and the slow depressor units, there is a pause in the activity of the extensor motor neurons. SRP1 is the most frequently observed pattern of coordinated motor activity in this recording (Fig4b). This substantiates previous results regarding the mesothoracic ganglion by Büschges et al. (1995). Note that not all depressor bursts are related to an SRP1, especially not those consisting of only the low-amplitude slow depressor units (Büschges et al. (1995)). Out of a total number of 1093 depressor bursts recorded from six mesothoracic preparations only 37.9% are accompanied by an SRP1 or SRP2. The phase analysis only displays the three preparations (*n* = 621) in which the extensor motor neuron activity is also recorded. However, the phase histograms corresponding to all six preparations can be seen in Fig S2 in the Supplementary Material. The mean distribution of protractor, retractor, and extensor spikes within the depressor cycle show distinct peaks and troughs (Fig 4c and Fig S2). The mean angles of spiking activity relative to the depressor cycle with a 90% confidence interval are calculated for each animal preparation and the average values for all animals are given in Table 2. Some important numbers in the table are also highlighted in the text. Fig 4c (All bursts) illustrates the overall increase in retractor activity (circular mean [90% CI]: 113.2° [111.5°,114.9°]) and decrease in protractor activity (circular mean [90% CI]: 359.8° [358.4°,1.2°]) during depressor bursts. The figure also highlights the synchronous inactivity in the extensor motor neurons (circular mean [90% CI]: 231.6° [230.4°,232.8°]) during depressor activity (circular mean [90% CI]: 55° [54.4°,55.5°]). Considering only cycles where an SRP1 occured (*n* = 130) the distributions appear to be similar, as exemplified by the similar circular means. However, the histograms display sharper peaks and the switch from protractor (circular mean [90% CI]: 4.6° [3.2°,5.9°]) to retractor activity (circular mean [90% CI]: 117.4° [115.6°,119.2°]) becomes more evident (Fig 4c, SRP1). In contrast, the last group of histograms show the opposite switch from retractor (circular mean [90% CI]: 19.4° [342.4°,56.5°]) to protractor activity (circular mean [90% CI]: 104.6° [94.3°,114.9°]) during ongoing depressor activity and extensor inactivity. This is indicative of an SRP2 (Fig 4c, SRP2).

**Table 2:** Mean values of the circular mean and the upper/lower 90% confidence intervals (C.I.) of three isolated mesothoracic ganglia preparations. The overall phase of the depressor and extensor activity does not change regardless of the type of analysis. Protractor and Retractor “SRP1”-values are similar to the “All bursts”-values and dissimilar to the “SRP2”-values.

| Activity | Analysis | Circ. Mean ($^\circ$) | 90% C.I. ($^\circ$) Upper | 90% C.I. ($^\circ$) Lower |
| --- | --- | --- | --- | --- |
| Depressor | All bursts | 55.0 | 55.5 | 54.4 |
|  | SRP1 | 56.1 | 56.8 | 55.5 |
|  | SRP2 | 47.1 | 49.9 | 44.4 |
| Protractor | All bursts | 359.8 | 1.2 | 358.4 |
|  | SRP1 | 4.6 | 5.9 | 3.2 |
|  | SRP2 | 104.6 | 114.9 | 94.3 |
| Retractor | All bursts | 113.2 | 114.9 | 111.5 |
|  | SRP1 | 117.4 | 119.2 | 115.6 |
|  | SRP2 | 19.4 | 56.5 | 342.4 |
| Extensor | All bursts | 231.6 | 232.8 | 230.4 |
|  | SRP1 | 235.9 | 237.4 | 234.5 |
|  | SRP2 | 224.0 | 233.3 | 214.7 |

**Figure 4:**
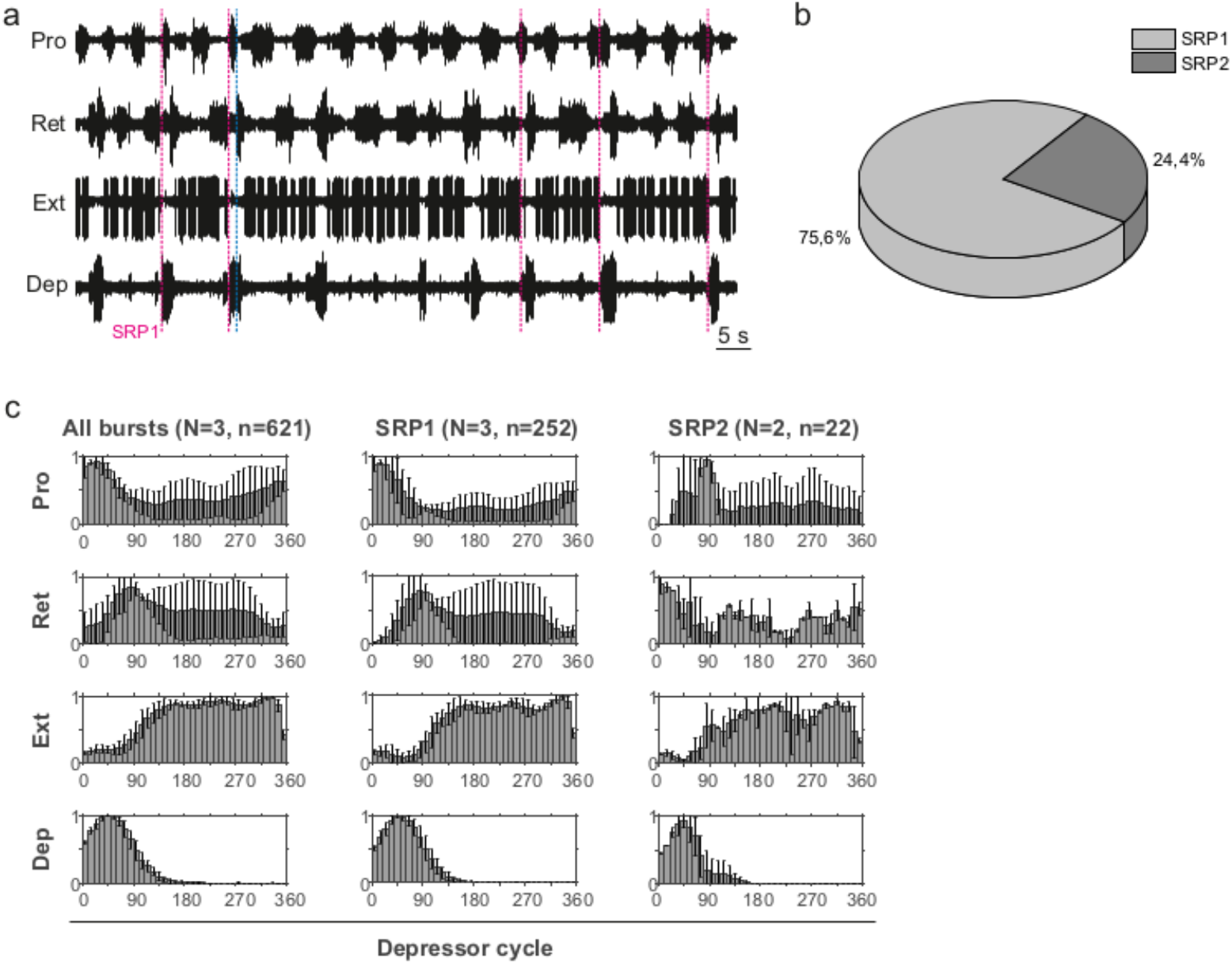
**a1**. Extracellular recording of the ipsilateral protractor (Pro), retractor (Ret), extensor (Ext), and depressor (Dep) motor neuron activity of an isolated (disconnected from other ganglia and deafferented) mesothoracic ganglion after bath application of 5*mM* pilocarpine. Motor activity becomes transiently coordinated when a switch from protractor to retractor activity occurs (blue dashed line) during an ongoing depressor burst and a pause in extensor activity (SRP1). Magenta dashed lines mark the depressor burst onsets during SRP1s. **a2**. Same preparation as in Fig 4a1. There is a switch from retractor to protractor activity (blue dashed line) during an ongoing depressor burst and a pause in extensor activity (SRP2). The orange dashed line marks the depressor burst onset during an SRP2. **b**. SRP1s occur more often than SRP2s in the recording in (a). **c**. Spike-phase histograms relative to the depressor cycle throughout the recording (All bursts) or during cycles where only an SRP1 or SRP2 occurs. In each histogram the mean (± STD) of each bin value among animal preparations is plotted. The y-axis represents average normalized number of spikes. “N” corresponds to the number of animal preparations and “n” to the number of depressor cycles.

Taken together, these results show that hemisegmental activity in the deafferented nerve cord can indeed be coordinated. Further revealing a higher occurrence of SRP1 in the isolated mesothoarcic ganglion. This represents a fictive transition from swing to stance phase during forward stepping.

#### 3.1.2 SRP2 occurs more often in the isolated meta- than in the meso-thoracic ganglion

Extracellular recordings of the ipsilateral motor activity after pharmacological activation with pilocarpine also reveal the occurrence of SRPs in the isolated and deafferented metathoracic ganglion (Fig 5a1 and Fig 5a2). SRP2s are denoted in Fig 5a1 with a dashed orange line and SRP1s with a dashed magenta line (Fig 5a2), always crossing through the onset of the respective depressor burst. Similar to the mesothoracic ganglion, from a total number of 890 depressor bursts recorded from six metathoracic preparations only 35.7% are accompanied by an SRP1 or SRP2. Extensor activity pauses each time there is combined activity of both the fast and slow depressor units (data not shown in Fig 5a). SRP2 occurs more often than SRP1 in this recording (Fig 5b). The distribution of protractor and retractor spikes throughout 752 depressor cycles show distinct peaks and troughs (Fig 5c (All bursts)). Again, the mean angles of spiking activity relative to the depressor cycle with a 90% confidence interval are calculated for each animal preparation and the average values for all animals are given in Table 3. Figure 5c (All bursts) illustrates the switch from retractor (circular mean [90% CI]: 239.3° [237.3°,241.4°]) to protractor activity (circular mean [90% CI]: 69.3° [68.8°,69.8°]) shortly after the onset of the depressor cycle (circular mean [90% CI]: 67.2° [66.5°,67.8°]), as denoted by the peak at approximately 0° in the retractor histogram. Considering only the cycles where an SRP1 occurs (*n* = 35), the initial switch from retractor to protractor activity is missing and only the opposite switch from protractor to retractor becomes evident at 40° to 45° (Fig 5c, SRP1). Here, the circular mean of the protractor (circular mean [90% CI]: 46° [45.3°,46.7°]) is smaller than the mean angle of the depressor (circular mean [90% CI]: 66.9° [65.8°,68°]). In the last group of histograms, based only on cycles where an SRP2 occurs, the initial switch from retractor to protractor activity during ongoing depressor activity reappears (Fig 5c, SRP2) and the circular mean of the protractor activity (circular mean [90% CI]: 80.8° [79.7°,81.9°]) is larger than the depressor mean (circular mean [90% CI]: 71.1° [69.1°,73.1°]). This is similar to the histograms corresponding to “All bursts”. The circular mean values point out the difference in the activity of the protractor and retractor between the deafferented preparations of the isolated meso- and meatathoracic ganglia. Therefore, coordination of hemisegmental activity in the deafferented metathoracic ganglion is qualitatively different compared to the mesothoracic ganglion and there is a higher occurence of SRP2s, representing a fictive transition from swing to stance phase during backward stepping.

**Table 3:**
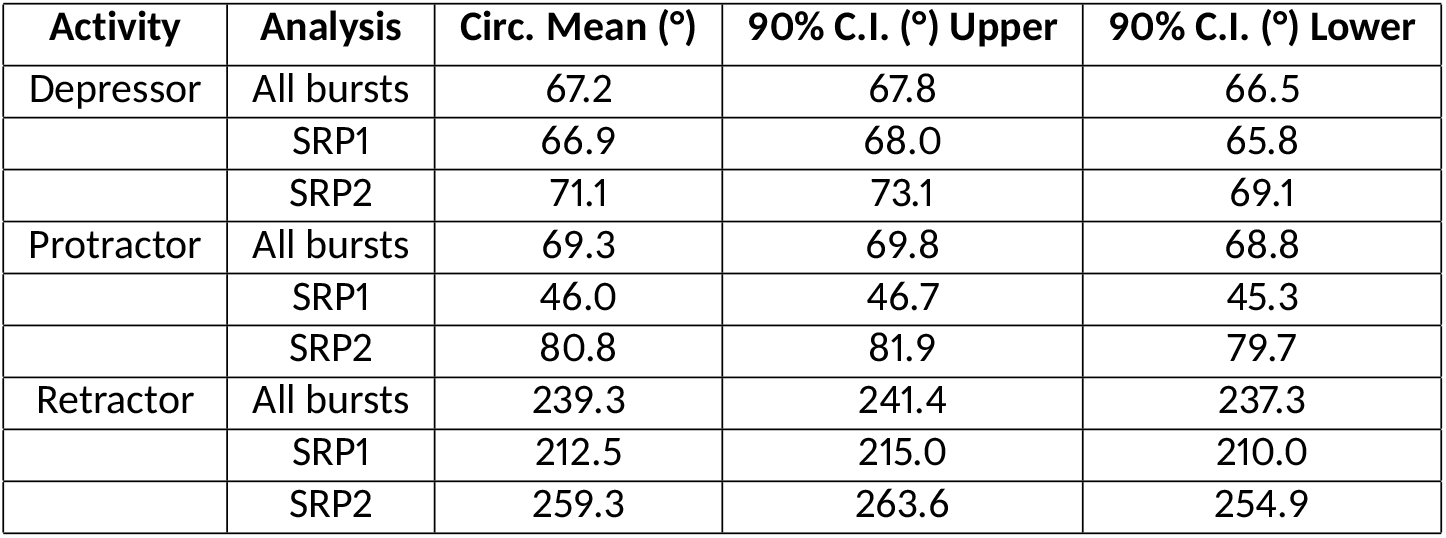
Mean values of the circular mean and the upper/lower 90% confidence intervals (C.I.) of five isolated metathoracic ganglia preparations. The overall phase of the depressor activity does not change regardless of the type of analysis. Protractor and Retractor “SRP2”-values are closer to the “All bursts”-values and dissimilar to the “SRP1”-values.

**Figure 5:**
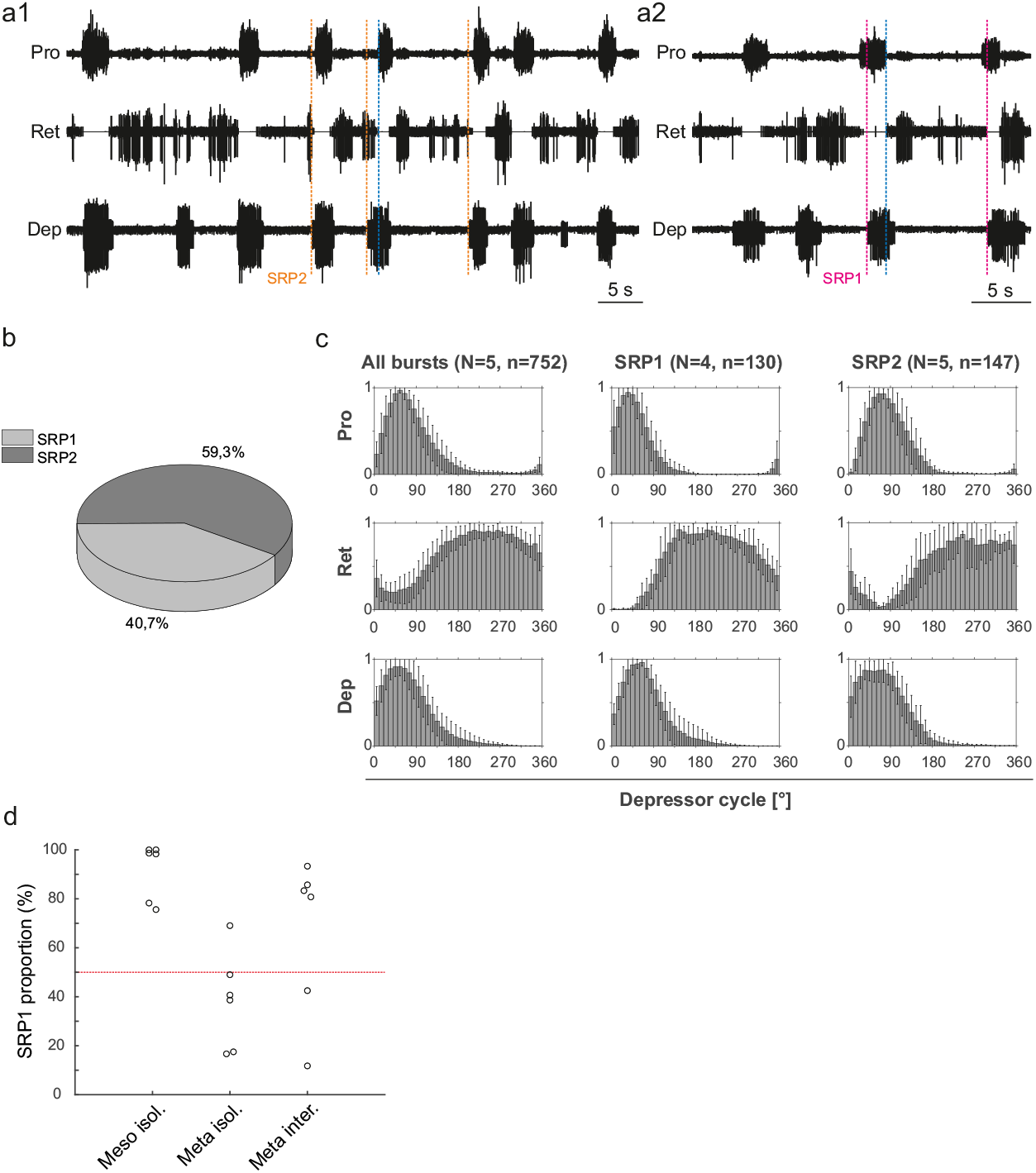
**a1**. Extracellular recording of the ipsilateral protractor (Pro), retractor (Ret), and depressor (Dep) motor neuron activity of an isolated metathoracic ganglion after bath application of 5*mM* pilocarpine. Motor activity becomes transiently coordinated when a switch from retractor to protractor activity occurs (blue dashed line) during an ongoing depressor burst (SRP2). Orange dashed lines mark the depressor burst onsets during SRP2s. **a2**. Same preparation as in Fig 5a1, magenta dashed lines mark the depressor burst onsets during SRP1s and the blue dashed line points out the switch from protractor to retractor activity. **b**. SRP2s occur more often than SRP1s in the recording in (a). **c**. Spike-phase histograms relative to the depressor cycle throughout the recording (All bursts) or during cycles where only an SRP1 or SRP2 occurs. In each histogram the mean (± STD) of each bin value among animal preparations is plotted. The y-axis represents average normalized number of spikes. “N” corresponds to the number of animal preparations and “n” to the number of depressor cycles. **d**. Proportion of SRP1s calculated from the sum of SRP1 and SRP2 patterns observed in recordings of the isolated meso- and metathoracic, and interconnected metathoracic ganglia preparations. There is an overall higher occurence of SRP1s in the isolated mesothoracic ganglion.

Fig 5d is based on data from different animal preparations. This plot reinforces the qualitative difference in ipsilateral coordination between meso- and metathoracic ganglia. The relative proportions of SRP1s and SRP2s are measured in the isolated meso- and metathoracic ganglia, and in the interconnected metathoracic ganglion ((*n* = 6) in each case). The proportion of SRP1s is significantly higher than 50% in the isolated mesothoracic ganglion (p=0.016, one-sample Wilcoxon signed rank test), whereas five out of six isolated metathoracic ganglion preparations show SRP1 proportions below 50% (Fig 5d). SRP1 proportions in the isolated and interconnected metathoracic ganglion do not significantly differ from 50% (p=0.219 amd p=0.437, respectively). However, four out of six interconnected metathoracic preparations show SRP1 proportions over 50%, implying a possible effect of intersegmental information on ipsilateral CPG coordination in the metathoracic ganglion (see Section 4). Taken together, our results indicate there is a higher frequency of SRP1s in the isolated meso-in contrast to a higher frequency of SRP2s in the isolated metathoracic ganglion.

### 3.2 Network Simulation

#### 3.2.1 Network topology promotes coordinated motor activity

The main result of the simulation is the network architecture itself. The configuration of a single joint pictured in Fig 2a is designed and confirmed to produce anti-phasic output based on the suggested architecture by the biological study from Yeldesbay and Daun (2020). This study informed our use of NSIs between the RGPs and MNPs, revealing a critical layer in the single joint architecture to allow precise control of output amplitude. The successful regulation of amplitude across frequencies is displayed in Fig 6.

**Figure 6:**
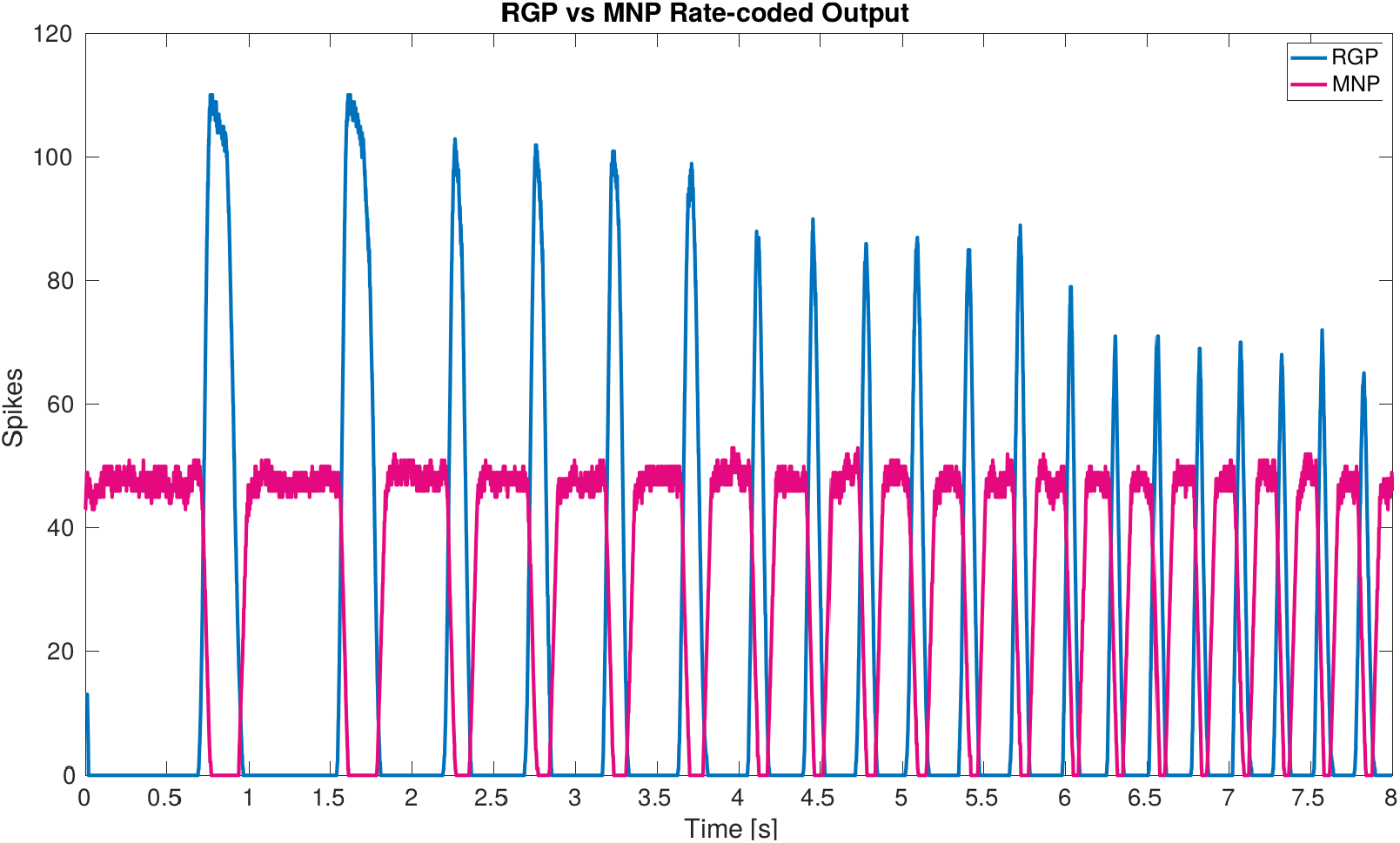
Plot of rate-coded output from the MNPs and the RGPs. The amplitude (amount of spikes) changes according to frequency from the RGPs but remains constant for the MNPs. This is achieved by using a constant bias current to achieve tonic spiking by the MNPs and using rhythmic inhibition from NSIs driven by the RGPs to create oscillations.

Furthermore, we find NSIs to be an effective method to loosely couple joints because they separate network dynamics. The addition of NSIs for intra-leg communication is inspired by the observations of Büschges (1995) indicating the role of NSIs during coordinated firing. In this way, our study provides corroborating evidence that NSIs may be important for coordination. Furthermore, our results indicate the insect may switch between different network architectures (communication pathways) based on walking direction. The architecture for the individual networks producing either forwards or backwards fictive step transitions is presented in Fig 2b and c.

#### 3.2.2 Simulated coordination is comparable to biological measurements

In order to confirm that the developed network architecture is biologically-plausible, its output is compared to biological measurements from a deafferented stick insect preparation. The comparison of network output to biological measurements starts with visual inspection to ensure the MNPs are bursting in the correct order to produce a swing to stance transition. Fig 7 shows the biological measurements on the left and the simulation results on the right for SRP1 and SRP2. As the measurements are at different timescales, the plots are normalized over the cycle period of each test respectively.

**Figure 7:**
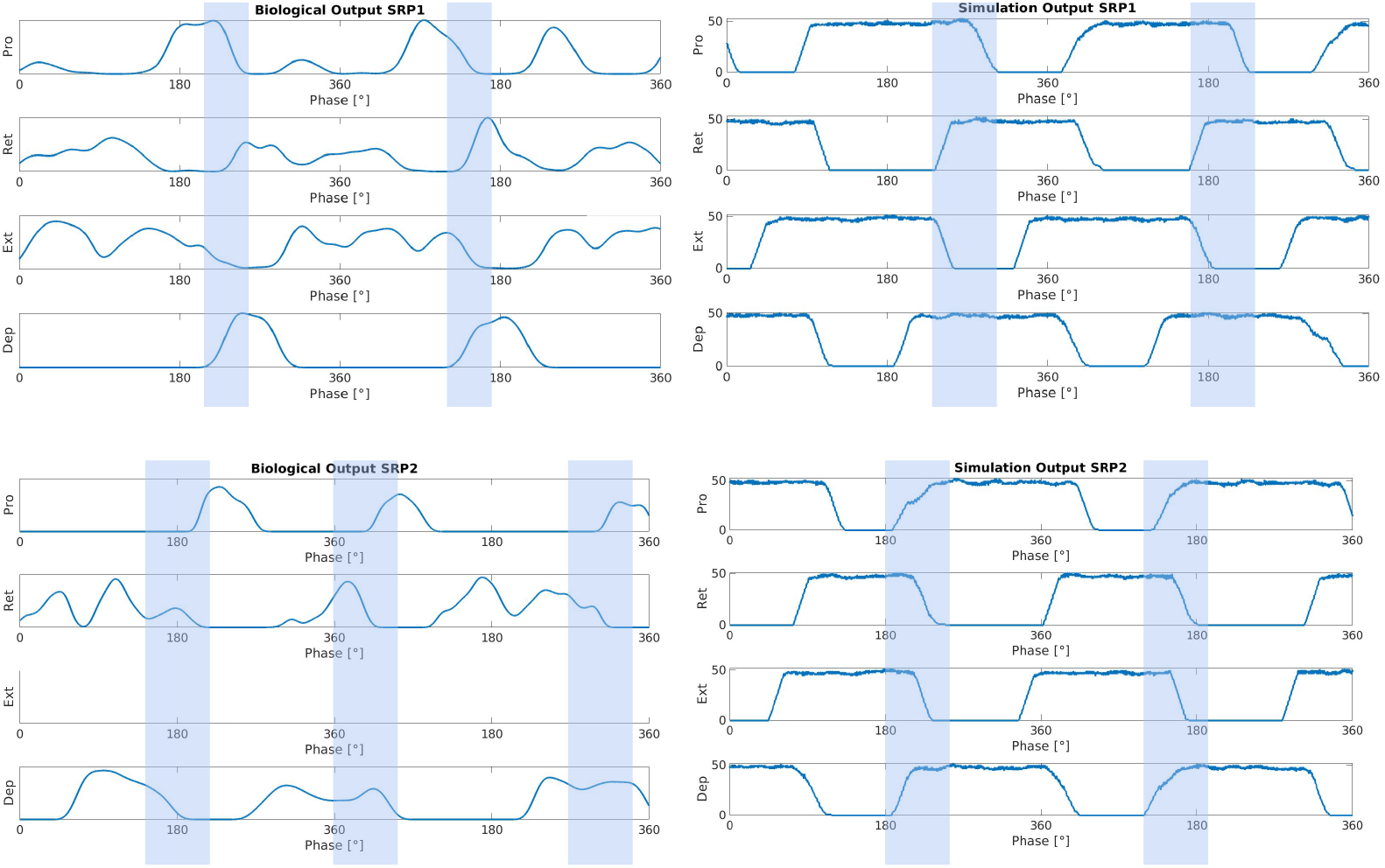
Side by side comparison of biological measurements (left) and output from the network simulation (right) during SRP1 (top) and SRP2 (bottom). The x-axis is the phase in degrees for all plots. The y-axis is normalized smoothed spiking activity for the biological output or number of spikes for the simulation output. The biological output for SRP1 is from the same animal as pictured in Fig 4a, also referred to as “Animal 1”. The simulation output also pictures “Simulated Animal 1”. The biological output for SRP2 is from the same animal as pictured in Fig 5a, also referred to as “Animal 4”. The simulation output also pictures “Simulated Animal 4”, which is the same as “Simulated Animal 3” because the two animals’ phase difference measurements for SRP2 are similar. During the depressor burst there is a switch from protraction to retraction in the SRP1 plots. Conversely, there is a switch from retraction to protraction during the depressor burst in the SRP2 plots. These transitions are highlighted within the blue boxes. Data was not available from the extensor during biological SRP2 recordings so this plot is empty but still pictured to keep consistency. The recordings from Animal 1 are from the isolated mesothoracic ganglion and Animal 4’s recordings are from the isolated metathoracic ganglion.

The figure confirms that during SRP1 there is switch from protraction to retraction during the depressor burst. Similarly, during SRP2 retraction ends and protraction begins during the depressor burst. This pattern is seen in both the biological and simulated results.

All tests of the network architecture are completed by switching from uncoordinated to coordinated firing after 2.5 seconds, allowing the network to initialize before switching to coordination. Four different phase difference ranges are evaluated, mimicking measurements from four individual animals (Animals 1-4) producing either SRP1 or SRP2 fictive step transitions. Each phase range is achieved at four different frequencies from 1− 4*Hz* to ensure results are validated across observed walking speeds of the stick insect (Graham and Cruse, 1981). The results confirm the developed architecture is able to produce a biological phase range when switching from uncoordinated to coordinated firing across the frequencies tested.

Fig 8 plots the calculated phase differences from the network simulation on top of the measured biological phase differences in motor activity from a single animal using the angular deviation to create the minimum and maximum values.

**Figure 8:**
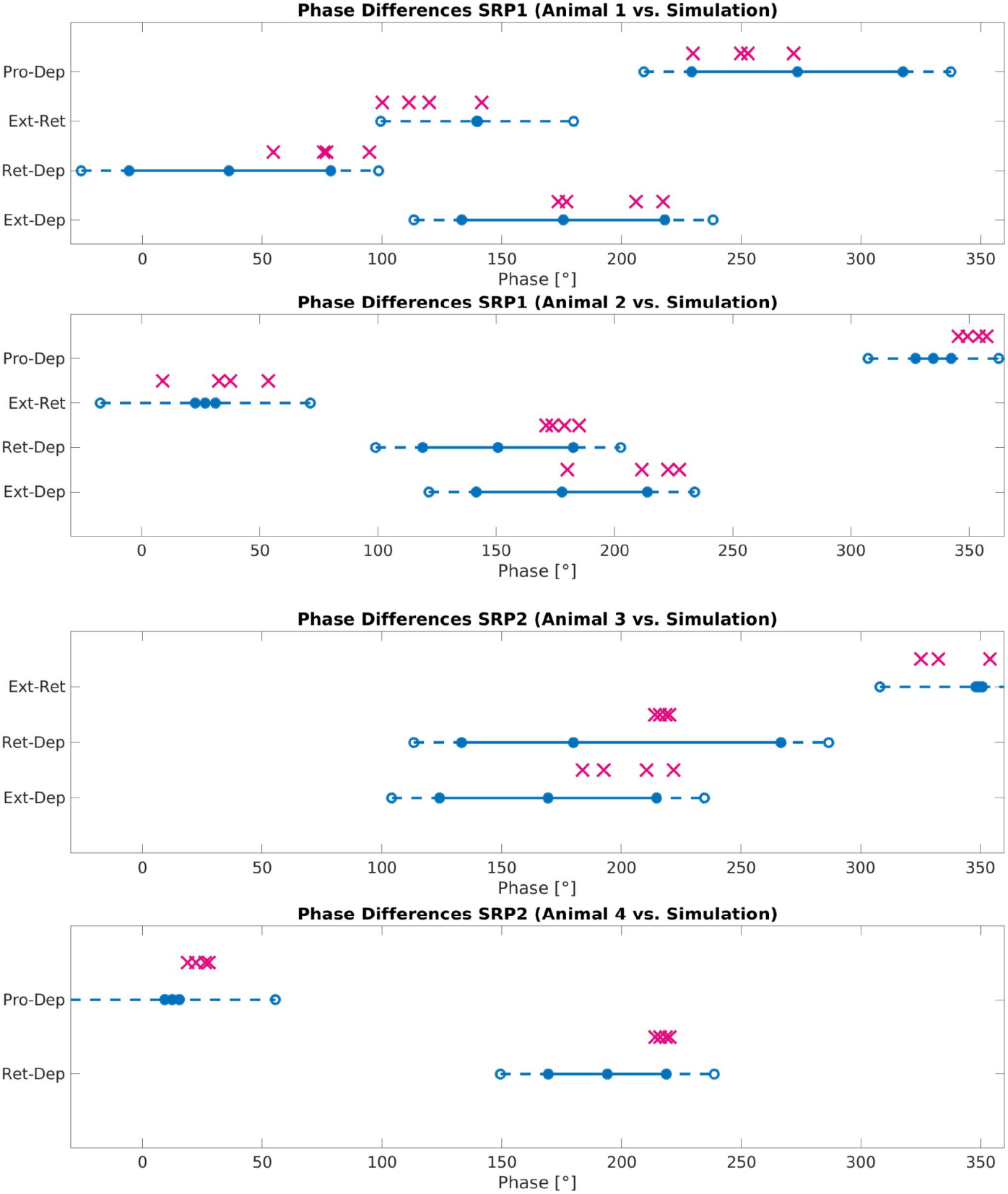
Phase difference between motor activity measured within deafferented samples (calculation method described by Fig 1) and in simulation (calculation method described by Fig 3 and Eq (11)). The filled circles and lines indicate the measured biological phase differences using the angular deviation from the mean to create a minimum and maximum. The empty circles and dashed lines indicate the buffer as described in Section 2.2.4. The ‘X’ marker indicates the phase difference recorded from simulation. There are 4 X’s per motor activity comparison reflecting each of the four frequency values tested. Each plot represents a single animal for a total of 4 different animals represented. The plots are labeled accordingly from Animal 1-4. Animal 1 is the same animal as pictured in Fig 4a, Animal 4 is the same animal presented with results in Fig 5a. The recordings from Animals 1 and 2 are from the isolated mesothoracic ganglion and from the isolated metathoracic ganglion for Animals 3 and 4.

Testing reveals that the network architecture must be altered to produce SRP1 and SRP2. The depressor population drives coordination during SRP1 whereas the levator population is used during SRP2. This difference is a result of manually testing multiple architectures, finding that the topology in Fig 2c is the only one capable of meeting biological phase difference measurements. However, regardless of the driving population, tuning the single parameter of synaptic delay from the ThC joint RGP to cNSI1 (see Fig 2b,c in the Methods section) is enough to match biologically measured phase differences in two different animals during both forward and backward walking. The exact synaptic delays from the RGP to cNSI1 used in this study can be found in Table 4.

**Table 4:** Synaptic delay from the CTr RGPs to cNSI1 producing biological phase differences per animal - Animal 1 (A1), Animal 2 (A2), Animal 3 (A3), Animal 4 (A4).

| Freq (Hz) | SRP1 A1 (ms) | SRP1 A2 (ms) | SRP2 A3 & 4(ms) |
| --- | --- | --- | --- |
| 1.3 | 90 | 350 | 70 |
| 2.2 | 70 | 210 | 30 |
| 3.2 | 50 | 120 | 5 |
| 4.0 | 30 | 90 | 239 |

The exact phase differences calculated from simulation output are also recorded in Tables S2, S3, and S4 in the Supplementary Material. As seen in Fig 8, the simulation is able to produce biologically-plausible phase ranges across the tested frequencies.

The system contains noise and investigation at different noise levels injected to the spiking populations indicates certain ideal levels depending on the individual animal’s coordination and walking direction. A standard deviation of 400*pA* is found to work for simulating Animal 1 to produce SRP1 while simulating Animal 2 needed 300*pA* of current noise to produce the correct phase differences. Phase ranges exhibited by two different animals (Animals 3 and 4) producing SRP2s could be reached in simulation using 400*pA* of current noise.

#### 3.2.3 Applying the simulated network on a robot leg shows stepping transitions

The output of the network is also tested on a physical robot simulator, CoppeliaSim (E. Rohmer, 2013), to visually confirm walking coordination on a robot leg. A link to the video can be found in the Supplementary Material (Video S1). Fig 9 shows a sequence of stepping behavior obtained from the video during coordinated firing.

**Figure 9:**
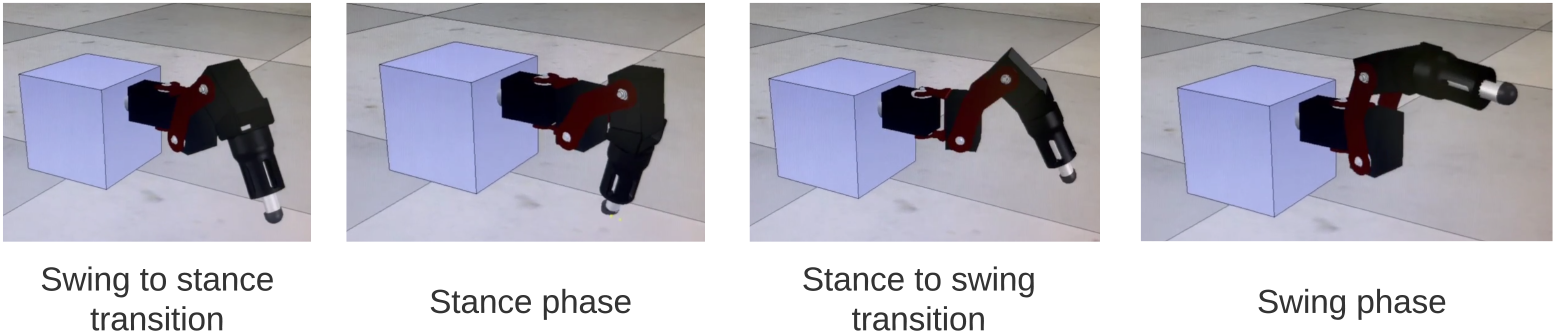
Stills from simulation of SRP1 using phase difference ranges measured from Animal 1 to create Simulated Animal 1. The stills reflect a full swing-stance step cycle.

The ability to control a single robot leg in simulation indicates the feasibility of eventually using this network with an additional interlimb coordination control mechanism (Aoi et al., 2017) to control a physical legged robot.

## 4 Discussion

### 4.1 Neurophysiology

Our study finds that SRPs occur not only in the meso- but also in the metathoracic ganglion. We substantiate the higher occurrence of SRP1s in the mesothoracic ganglion, as previously described in Büschges et al. (1995), and we report a higher occurence of SRP2s in the metathoracic ganglion. As SRP2 represents a fictive transition from swing to stance phase during backward stepping, it is plausible to assume there is an inherent tendency for forward walking in isolated meso- and for backward walking in isolated metathoracic ganglia. This result is in line with findings by Bässler et al. (1985), reporting that the hind legs of the stick insect perform backward stepping when they do not receive input from anterior legs, indicating an inherent backward direction of movement for the metathorax. Interestingly, this segment specificity in relation to stepping direction can also be demonstrated after sensory stimulation. Load increase at the level of the mesothorax after pilocarpine application results in retractor activation, whereas at the level of the metathorax the protractor is activated, pointing towards activation of forward and backward stepping respectively (Akay et al., 2007). Based on the mentioned studies and our results, we conclude that the meso- and metathoracic networks for stepping may rely on different architectures.

We also show the proportion of SRP1s is above 50% in four out of six preparations of the interconnected metathoracic gangion. Future studies should increase the sample size and verify our observation, which implies that intersegmental information descending from the mesothoracic ganglion may affect coordination of motor activity at the metathoracic ganglion, ultimately changing the fictive backward stepping phase transition (SRP2) to forward (SRP1). A similar change in the activity is also observed *in vivo*, as the inherent backward direction of the metathorax changes to forward when all other legs are stepping (Bässler et al., 1985). Additionally, it has been found that intersegmental information from the middle legs is necessary to produce regular stepping in the hind legs in freely walking animals (Grabowska et al., 2012). Another notable phenomenon is that left-right coordination of motor activity is modified in the metathoracic ganglion when it is interconnected to the mesothoracic ganglion or the rest of the nerve cord in deafferented stick insect and locust preparations (Mantziaris et al., 2017, Knebel et al., 2017). Considering all of the above, we present evidence for segment specificity in the walking system of the stick insect. The difference in the underlying neural mechanisms among segments and the functional significance of our results will be the focus of our future experiments.

### 4.2 Network Simulation

Our study reveals that the addition of 2 cNSIs is enough to coordinate joints and produce fictive step transitions. However, the synaptic connections to these cNSIs must be altered depending on the direction of walking. SRP1 requires the depressor, retractor, and flexor to fire together, therefore, coordination is created through coupling their respective RGPs directly (see Fig 2b in the Methods section). SRP2 needs the levator, retractor, and extensor to fire at the same time so the architecture switches to driving coordination from the levator RGP and exciting the retractor and extensor RGPs (see Fig 2c in the Methods section). This indicates that there is a distinct difference in synaptic connectivity during forwards and backwards step transitions and could explain why the metathoracic ganglion produces mostly SRP2s when disconnected from the other ganglia. This finding is similar to the biological outcome from Büschges (1995) showing that exciting certain NSIs increased the likelihood of one type of SRP versus another. The replication of this phenomenon in simulation by means of altering network connectivity, points to a partial role of network architecture in stepping patterns.

Fig 7 shows that the burst duration in simulation is much longer than in biological experiments. Therefore, there is more overlap when the MNPs are spiking. This does not affect the phase difference calculations since these only account for onset in spiking. Based on the visual observation of the robot leg in simulation, the transitions can be subjectively judged as acceptable even though there is significantly more spiking overlap. However, this must be further investigated when building upon the suggested network. It should also be noted that the phase difference calculation is from onset of spiking when evaluating the simulation data (Fig 3) as opposed to an average of all spikes in a burst when looking at biological data (Fig 1). We assert this is comparable because burst duration remains stable at any given frequency ((Büschges et al., 1995)). Therefore, comparing the average versus the onset will produce the same phase difference because the mean of the burst time remains equidistant to the onset of the burst time.

Our developed network architecture using both spiking and non-spiking neurons to regulate output from the MNPs should be considered for use in other research, expanding on the commonly used half-center oscillator (Marder and Bucher, 2001, Bidaye et al., 2018) architecture associated with CPGs. The addition of NSIs has several advantages including separating the dynamics of the system and increased control over input to spiking populations. As seen in Strohmer et al. (2021), the output amplitude of the MNPs changes with frequency due to the use of a constant time window when rate-coding spikes. This variance is removed by adding buffer NSIs between the RGPs and MNPs (see Fig 6). The affect of the excitatory spikes from the RGPs is limited by the synaptic weight to the NSIs and neuronal dynamics of the NSIs themselves. The change in membrane potential of the NSI determines the inhibitory current sent to the MNPs acting as a way to smooth noisy spiking input. In addition to incoming spikes, current injection also affects an NSI’s membrane potential. The amount of current and whether it is excitatory or inhibitory, regulates the membrane potential fluctuation of an NSI. The ability of NSI’s to receive spikes or current and translate them into a predictable membrane potential fluctuation makes them useful for connecting neural network architectures or conveying sensory information to a neural network.

The rhythmicity of network output is driven by the RGP neurons. Coarse frequency modulation is achieved through changing the voltage threshold potential of the neurons in the RGPs. This relationship was found by Strohmer et al. (2021) where a change of approximately 1*Hz* is achieved by manipulating the voltage threshold potential by 1*mV*. This relationship only holds until a voltage threshold minimum of *V*_*th*_ = − 57*mV* due to the other chosen neuronal characteristics. This *V*_*th*_ still produces an output of approximately 2*Hz* when using the original study’s suggested excitatory current injection of 500*pA* to the RGPs (Strohmer et al., 2021). Therefore, further tuning is required to reduce the frequency to the desired 1*Hz* minimum. During testing, fine frequency modulation is achieved by adjusting the level of current injection to the RGP. Reducing the excitatory current to 20*pA* is able to reduce the frequency to approximately 1.2*Hz* which is accepted to be close enough to the desired minimum. The capacitance of the RGPs also needs to be increased to 400*pF* to smooth the output at such low frequencies.

Phase difference is manipulated through synaptic delays from spiking populations to NSIs. The delay to cNSI1 (see Fig 2b,c) can be used to adjust the phase based on frequency. This tuning parameter is able to replicate the phase differences measured on animals displaying motor activity coordination as far as 114° apart (Table S1 in the Supplementary Material), exhibiting the sensitivity of the system to this variable. Video S1 (in the Supplementary Material) of the simulation confirms that the motor activity coordination between biological animals is significantly varied. Visual observation shows the Simulated Animal 1 has a more intuitive phase difference during swing to stance transition which more closely resembles forward walking in a live animal. As noted in the results, the optimal amount of noise to produce the correct motor activity coordination in simulation for forward and backward swing to stance transitions per animal differed. This is considered acceptable since noise is by definition not constant.

The network simulation phase differences presented in Fig 8 are an average over all calculations occurring after coordinated firing is initiated. However, SRPs are spontaneous and not cyclic so further testing must be done to see if the network can produce singular instances of coordinated firing. Evaluating individual phase differences between motor activity shows that it can take between 0.14 − 2.85*s* before the compared phase of the motor activity is within the accepted range as compared to biological measurements (see Tables S2, S3, S4 in the Supplementary Material). This difference in “initialization time” could be caused by the significant amount of noise in the system and should also be further evaluated.

The main finding that two NSIs can be used to couple the leg joints to create coordinated firing allows for either descending signals or sensory feedback to be easily added to the system. NSIs can receive spikes or current injection, making them well suited to connect with spiking sensory neurons or translate analog sensory information. The identification of analogous interneurons in the biological system and possible sensory organs affecting their activity should be the focus of future research.

## Author Contributions

BS and CM developed the main idea of the paper and contributed equally to this work. BS researched the biological neural network concepts, formulated a theory, and applied it in simulation. CM and DK performed the biological experiments and CM analyzed the associated data. The manuscript was written by BS and CM with support from LBL, AB, and PM. AB supervised the project. LBL and PM discussed the results. LBL and AB provided the funding. All authors contributed to the article and approved the submitted version.

## Acknowledgements

The authors would like to thank Mathias Thor for providing access to the MORF robot leg in the simulation environment.

This paper is available on the BioRxiv pre-print server.

