## Supplementary Material for "Network architecture producing swing to stance transitions in an insect walking system"

bioRχiv
THE PREPRINT SERVER FOR BIOLOGY

### Network architecture producing swing to stance transitions in an insect walking system

Beck Strohmer[1,*,†], Charalampos Mantziaris[2,†], Demos Kynigopoulos[3], Poramate Manoonpong[1], Leon Bonde Larsen[1], and Ansgar Büschges[2]

[1]SDU Biorobotics, Maersk McKinney Moller Institute, University of Southern Denmark
[2]University of Cologne, Department of Animal Physiology, Zoological Institute, Cologne, Germany
[3]The Cyprus Institute of Neurology and Genetics, Cyprus School of Molecular Medicine, Agios Dometios, Cyprus
[†]These authors contributed equally to this work.

#### Supplementary Material

**Video S1.** Video link showing stepping in simulation.

Table S1: Phase differences for Animals 1-4 calculated by subtracting the circular mean values. For example, if Ext-Dep is compared, the circular mean of the Dep motor activity is subtracted from the Ext. The fields marked "N/A" indicate where data was not available. This table does not show the angular deviation that creates the minimum and maximum biological range.

| Animal | Ext-Dep () | Ret-Dep () | Ext-Ret () | Pro-Dep () |
|---|---|---|---|---|
| 1 (SRP1) | 175.76 | 36.05 | 139.72 | 273.48 |
| 2 (SRP1) | 177.72 | 150.74 | 26.98 | 334.86 |
| 3 (SRP2) | 169.43 | 179.95 | 349.48 | N/A |
| 4 (SRP2) | N/A | 194.08 | N/A | 12.41 |

Table S2: SRP1 phase differences per frequency for Simulated Animal 1. Init (initialization) time refers to the amount of time from activating the cNSIs until the first recorded phase difference within biological measurements.

| Freq (Hz) | Ext-Dep () | Ret-Dep () | Ext-Ret () | Pro-Dep () | Init Time (s) |
|---|---|---|---|---|---|
| 1.25 | 173.63 | 54.71 | 119.77 | 229.79 | 1.26 |
| 2.17 | 177.03 | 76.83 | 100.20 | 252.86 | 2.85 |
| 3.21 | 206.05 | 94.77 | 111.28 | 271.96 | 0.15 |
| 4.04 | 217.30 | 75.74 | 141.58 | 249.83 | 0.14 |

Table S3: SRP1 phase differences per frequency for Simulated Animal 2. Init (initialization) time refers to the amount of time from activating the cNSIs until the first recorded phase difference within biological measurements.

| Freq (Hz) | Ext-Dep () | Ret-Dep () | Ext-Ret () | Pro-Dep () | Init Time (s) |
|---|---|---|---|---|---|
| 1.24 | 180.03 | 171.04 | 9.00 | 345.37 | 1.38 |
| 2.15 | 222.55 | 184.96 | 37.59 | 353.97 | 0.65 |
| 3.17 | 227.46 | 173.76 | 53.70 | 357.31 | 2.01 |
| 3.97 | 211.63 | 178.85 | 32.71 | 349.34 | 0.21 |

Table S4: SRP2 phase differences per frequency for Simulated Animals 3 and 4. Init (initialization) time refers to the amount of time from activating the cNSIs until the first recorded phase difference within biological measurements.

| Freq (Hz) | Ext-Dep () | Ret-Dep () | Ext-Ret () | Pro-Dep () | Init Time (s) |
|---|---|---|---|---|---|
| 1.25 | 183.81 | 218.59 | 325.22 | 18.96 | 0.47 |
| 2.19 | 192.65 | 220.16 | 332.49 | 22.36 | 0.54 |
| 3.19 | 210.57 | 216.08 | 354.07 | 27.66 | 0.15 |
| 3.99 | 221.79 | 213.97 | 7.83 | 26.23 | 0.14 |

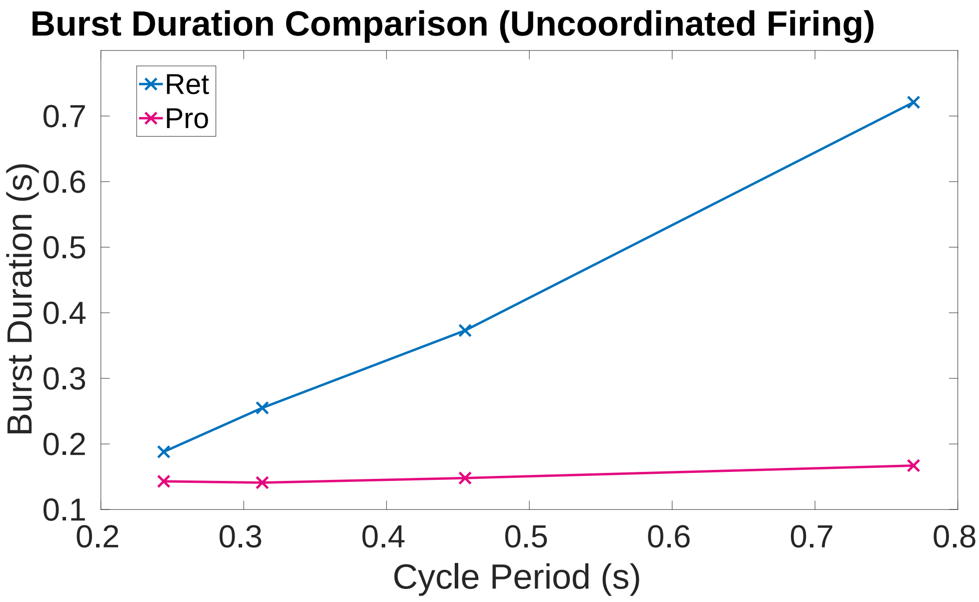


Figure S1: Plot of burst duration for the retractor (ret) and protractor (pro) MNPs at tested cycle periods. The ret MNP burst duration increases in correlation with the cycle period. The pro MNP burst duration is confirmed to be independent of cycle period.

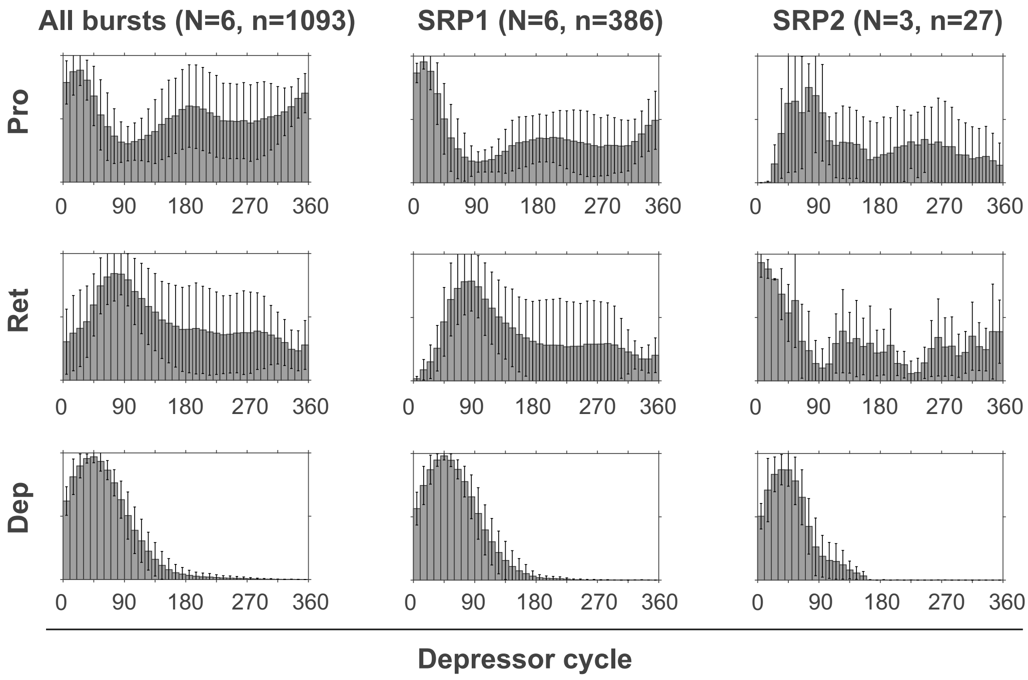


Figure S2: Data analysed from six isolated mesothoracic ganglion preparations (three of which are included in Figure 4). Spike-phase histograms relative to the depressor cycle throughout the recording (All bursts) or during cycles where only an SRP1 or SRP2 occurs. In each histogram the mean ($\pm$ STD) of each bin value among animal preparations is plotted. The y-axis represents average normalized number of spikes. "N" corresponds to the number of animal preparations and "n" to the number of depressor cycles.
